# Extracting Parsimonious Quantitative Predictors of Biological Effectiveness from ‘First-Principles’ Radiobiology: Application to the Mixed-Quality Problem

**DOI:** 10.64898/2026.05.02.722446

**Authors:** Tahir I. Yusufaly, Mark K. Transtrum, Livia Huang, Seyed A. Sabok-Sayr, George Sgouros, Robert F. Hobbs, Xun Jia

## Abstract

Developing parsimonious, mechanism-aware quantitative models that predict how biological effectiveness changes with different modifiers remains, in general, an unsolved problem. Advances in radiobiological research have created a large knowledge base of ‘first-principles’ mechanistic models of radiation response that, in principle, could accurately predict radiosensitivity across different experimental and clinical conditions. However, in practice these mechanistic models come with an overabundance of parameters, the majority of which are practically unidentifiable and, moreover, likely unnecessary if one simply wishes to predict how radiosensitivity changes for some specific modifier of interest. Nevertheless, determining which few details in the full mechanistic model are relevant for a given purpose, as well as how to remove any other extraneous details, remains a highly non-trivial task. In this study, we demonstrate the potential of model reduction, starting from a detailed mechanistic description, as a systematic strategy for deriving parsimonious, experimentally falsifiable radiobiological descriptors. As a proof-of-concept demonstration, we apply the Manifold Boundary Approximation Method (MBAM) to a Mechanistic Model of DNA Repair and Survival (MEDRAS), for the problem of cell survival prediction following an acute exposure. Our findings reveal that the complete MEDRAS model for an arbitrary mixed-quality exposure can be structurally simplified to a reduced three-parameter model for an effective uniform-quality, named MEDRAS-LPL. Additional MBAM analysis on MEDRAS-LPL identifies two boundaries in parameter space, corresponding to sparsely ionizing and densely ionizing radiation. Mapping of MEDRAS-LPL parameter space on to effective LQ space further demonstrates that parameters close to the sparsely ionizing boundary line up with expectations from the theory of dual radiation, while parameters close to the densely ionizing boundary line up with expectations from a purely linear model based on a target-theory description. Moreover, our formalism predicts enhanced synergistic interactions between sparsely ionizing and densely ionizing radiation beyond the Zaider Rossi model (ZRM) paradigm, in line with empirical observations. The results highlight the potential for using reduced-order models not only for predictive applications but also for generating novel hypotheses that can inform future experimental designs and optimization strategies in radiobiology.

## 1. Introduction

Modern clinical radiobiology is centered around the linear-quadratic (LQ) model (1),

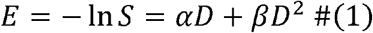

Here, *E* is the biological effect, which we define as the negative logarithm of the cell survival fraction *S. E* is a function of absorbed dose *D* and consists of two contributions. The linear term, parameterized by *α*, characterizes the exponential decrease in cell survival at low doses. The quadratic term, parameterized by *β*, is responsible for the ‘shouldering’ of the curve at higher doses. Despite its simplicity, at an empirical level the LQ model is remarkably robust. The biological effectiveness of radiation is modified by many possible factors (2,3), such as radiation quality (4), dose rate (5,6), intrinsic cellular phenotype (7,8), or local oxygenation (9,10). However, for a fixed set value of modifiers, the shape of the *S*(*D*) curve is almost universally observed to display an LQ dependence for low and intermediate doses per fraction (although extrapolation of the LQ at high doses per fraction remains debatable (11,12)), with the cumulative effect of the possible modifiers encapsulated in the specific values of the fit parameters *α* and *β*.

The parsimony of the LQ model makes it tremendously useful in radiation oncology. It serves as a general framework for comparing biological effectiveness across different scenarios. As a common example, a clinician with experience doing treatment planning under a standard fractionation schedule may wish to extrapolate that experience to an alternative schedule, or to a protracted low dose rate exposure due to continuous decay of a radionuclide. Such extrapolation can be performed straightforwardly by keeping the linear term α fixed and modifying the quadratic term *β* with a dose rate factor *G* (13) to calculate the biological effective dose (BED) (5,6).

In principle, such a strategy should be generalizable across a wide range of scenarios, such that if we have some arbitrary modifier of biological effectiveness *X*, the calibrated biological dose can be estimated based on the *X*-dependence of the LQ parameters, *α*(*X*) and *β*(*X*). Unfortunately, there are few situations in which we can explicitly calculate how the LQ parameters change as a function of *X*. In practice, therefore, these parameters must usually be separately measured for individual values of *X*, with limited transferability outside of rather specific conditions (14,15).

Occasionally, it is possible to empirically fit *α*(*X*) and *β*(*X*) data to some ‘black box’ function that is good at *interpolating* across measured *X* values. However, such efforts usually fall short of being able to *extrapolate* to *X* values outside the range that is measured. To be capable of such extrapolation, the functions *α*(*X*) and *β*(*X*) must, in practice, also be endowed with some kind of underlying mechanistic interpretability that enables them to make falsifiable predictions in truly novel situations. In the case of variable dose rates, for example, the *G* factor has an underlying biophysical interpretation in terms of the incomplete repair kinetics of sublethal DNA damage (13,16,17). This interpretability allows for a more systematic, targeted exploration of the space of possible time-dependent 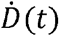, beyond trial-and-error random searching of different fractions, radionuclide half-lives, etc….

Unfortunately, the example of dose rate is something of an outlier, and for more general modifiers *X*, we typically lack a rational strategy for efficiently mapping the landscape of *α*(*X*) and *β*(*X*). The development of such a strategy, therefore, could offer the potential to dramatically expand the scope of BED modeling and estimation, in a way that could eventually allow for more flexible and adaptive treatment planning.

A promising starting point towards this goal are ‘first-principles’ Monte Carlo simulations of the underlying radiobiological mechanisms (18-21). While a *quantitative* mapping from a given modifier *X* to the corresponding *α*(*X*) and *β*(*X*) is usually not known, it is often the case that we have a reasonably correct *qualitative* understanding of the ways in which these modifiers couple to the underlying intracellular pathways of radiation response. The development of software such as Geant4-DNA (22) and its extension to TOPAS-nBIO (23) today allows for detailed track structure simulations of the radiation chemistry of water and the induction of simple and complex patterns of DNA damage at base-pair resolution, for a variety of cellular and nuclear geometries at various levels of realism. The outputs of these damage simulations can, in turn, be paired with models of the DNA damage response (24,25) and the resulting induction of cell death via a combination of early apoptosis and late mitotic catastrophe. Results from mechanistic simulations have been shown capable of qualitatively reproducing expected behaviors and trends across a variety of clinical conditions (26-29).

However, translating these insights from mechanistic, literature-derived models and simulations into practical tools for day-to-day clinical practice is challenging. The reasons for this are multifaceted, but arguably the most important factor is the overabundance of model parameters, the vast majority of which are practically unidentifiable experimentally or clinically. Indeed, recent studies (30,31) have shown that the predicted DNA damage yields and biological endpoints can vary significantly with reasonable variations in the mechanistic parameters. Thus, while *in principle* mechanistically parameterized simulations could offer a more fundamental and generalizable description of radiobiology than ‘black-box’ measurements of LQ phenomenology, *in practice* they are plagued by prohibitively large uncertainties.

Moreover, there is also a growing realization that, while first-principles Monte Carlo radiobiology is universal, it is also in most situations overkill (32). Indeed, the example of the G-factor shows that biophysical knowledge of how a specific modifier (dose rate) influences biological effectiveness can be incorporated in a way that is consistent with the underlying mechanistic knowledge, whilst simultaneously honing out only those details that are most essential for calibrating between fractionated and protracted exposures. But as mentioned above, dose rate is something of an anomaly, and in general, the identification of parsimonious, mechanistically predictive adjustments to the LQ parameters for arbitrary modifiers *X* is not yet possible.

Nevertheless, the progress in mechanistic radiobiological knowledge and Monte Carlo simulations can still be useful towards this goal. Suppose we have a reasonably accurate qualitative understanding of how a modifier couples to the underlying radiation response mechanisms, at a level that allows us to explicitly incorporate it into first-principles radiobiological modeling and calculations even if we don’t know the exact values of all the parameters. Then, it might be possible to start from this multi-parameter description and hone out the select few parameters-of-interest that are needed for predicting how changes in *X* map to changes in the effective phenomenological LQ survival parameters *α* and *β*. However, the task of model reduction, or deciding which parameters in the original mechanistic model are relevant to a given application and how to remove the extraneous irrelevant ones, is in general non-trivial.

Over the years, various approaches to model reduction across different research communities have emerged (33–36) to address the problem of finding the ‘right’ approximations for a given context. One technique that has recently proven fruitful in systems biology and pharmacology applications is the Manifold Boundary Approximation Method (MBAM (37)), which is built around the idea that the geometry of parameter space in complex multi-parameter models can be organized into ‘stiff’ and ‘sloppy’ directions (38–40). Stiff directions correspond to combinations of parameters that, when changed, most significantly impact a predicted experimental output, such as dose-response or survival fraction. Sloppy directions, on the other hand, correspond to parameter combinations that have relatively little influence on predicted input-output relationships. The removal of the sloppy directions corresponds to a series of physically interpretable approximations that allow the reduced model to retain mechanistic interpretability. MBAM has been applied to a range of systems biology problems, an early example (41) being the EGFR signaling pathway. These reduced models show promise for quantitative systems pharmacology (42) via fit-for-purpose explainable AI that could accelerate drug development and approval.

In this work, we extend the application of MBAM to the realm of radiobiology, specifically the task of model-based extrapolation of how effective LQ parameters vary across possible values of effectiveness modifiers *X*. Specifically, we demonstrate our approach in the context of the problem of mixed-quality radiation response (43), illustrated in **Figure 1**: given measured LQ parameters for two or more different types of radiation, with varying degrees of ionization density per unit dose, can we predict how the effective LQ curve will vary if we mix the different radiation qualities together in some proportion? In addition, we explore whether such model-based inference can guide us in the search for ‘optimal’ cocktails of qualities that maximize the potency of cell kill. Our results for the mixed-quality problem in this paper lay a more general foundation for going beyond trial-and-error empirical exploration of LQ radiosensitivity, towards a paradigm of mechanism-guided discovery and design.

**Figure 1:**
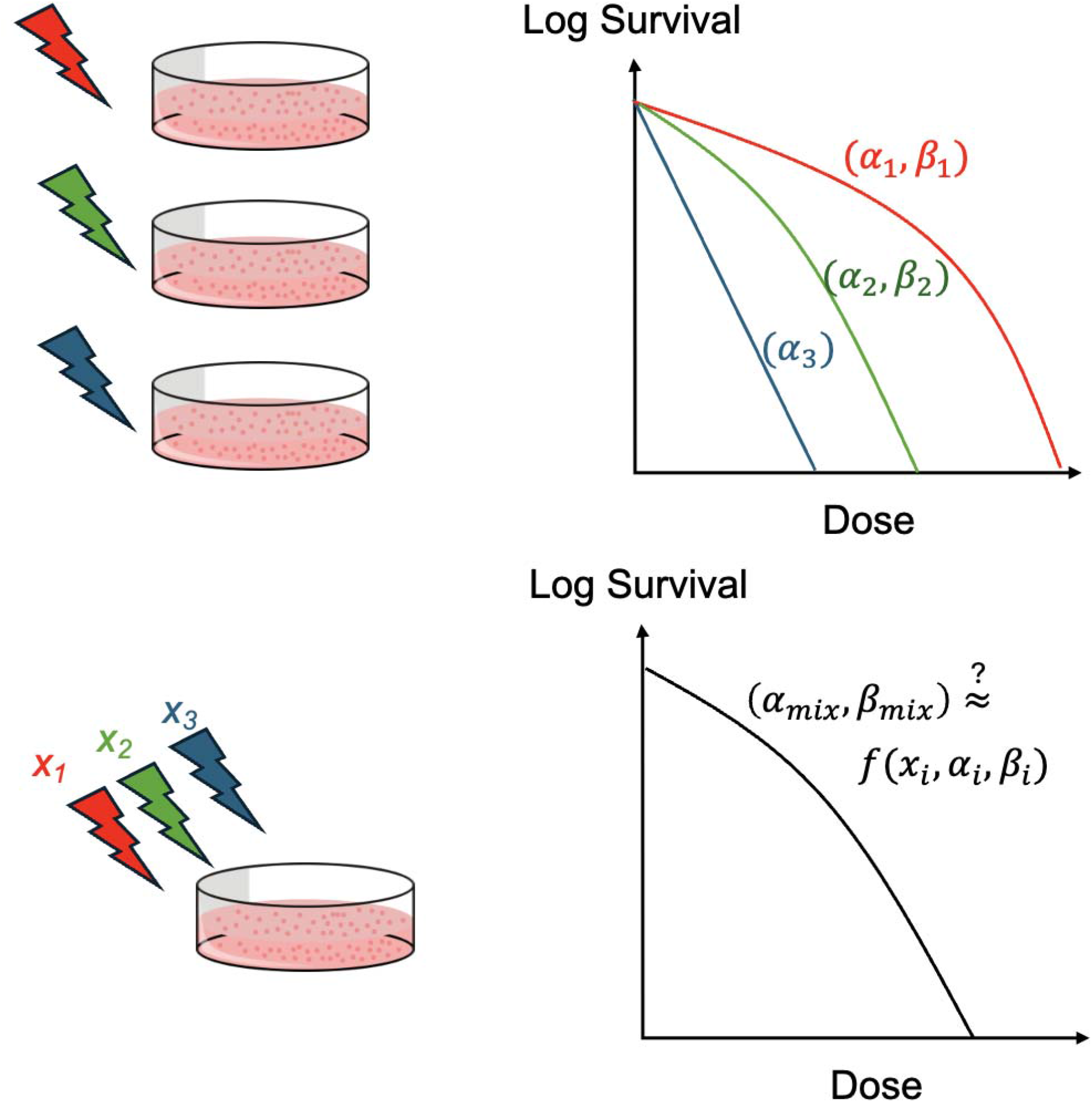
The radiobiological task of predicting survival curves for a mixed beam from the survival curves of the components remains challenging, particularly when the beams include a mix of sparsely ionizing (red) and densely ionizing (blue) qualities.

## II. Background

In this section, we introduce the necessary background knowledge to appreciate the results of this paper. We begin in subsection II.A. by reviewing the clinical context and importance of the mixed-quality radiation response problem, and the challenge of understanding how interactions between sparsely and densely ionizing radiations modulate effective LQ parameters. In subsection II.B., we introduce the basics of Mechanistic DNA Repair and Survival (MEDRAS (25)), a mechanistic radiobiological model that looks to incorporate state-of-the-art knowledge of how spatial damage distributions from different types of radiation interact with DNA damage response pathways to modulate repair or mis-repair.

### A. Mixed Quality Problem and Inadequacies of ZRM

The mixed quality radiation response problem, which involves the biological effects of exposure to different types and energies of radiation, presents several significant challenges in both clinical oncology and radiological protection. The most common approach to analyzing mixtures of radiation types is the Zaider-Rossi Model (ZRM) (44,45) based on the theory of dual radiation action (46,47), which we describe below.

For a single radiation type *i* delivered at dose *D*_*i*_, the standard LQ survival is

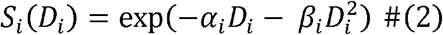

The theory of dual radiation action assumes that we can associate each term in the LQ formula with one of two classes of lethal lesions: the linear term *α* is associated with single-event lethal lesions, while the quadratic term *β* corresponds to pairwise interactions between sublethal lesions.

Now consider a mixture of radiation types *i* = *1*, …, *N*, each delivering dose *D*_*i*_,. Define the total dose *D* = *∑*_*i*_ *D*_*i*_ and dose fractions *f*_*i*_= *D*_*i*_ /*D* (so*∑*_*i*_ *f*_*i*_ = 1). We aim to find effective *α*_*mix*_ and *β*_*mix*_such that

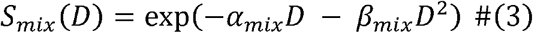

The ZRM treats single-event lesions as additive across radiation types. Meanwhile, the number of pairwise interactions between sublethal lesions is quadratically proportional to the total number of sublethal lesions, irrespective of the type of radiation the lesion originates from. These postulates result in the prescription

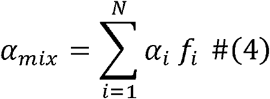

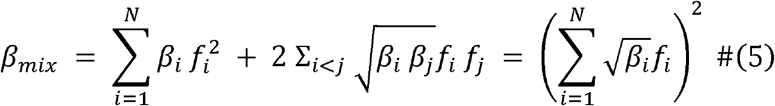

The mixed quadratic contribution includes self-quadratic terms (i=j) and cross terms (i≠j and the second equality follows from completing the square. These equations can be further generalized beyond simultaneous acute deliveries to account for more protracted and temporally structured exposures (48).

The ZRM approach is most accurate when the different types of radiation in the mixture are of similar quality (e.g., all sparsely or all densely ionizing). However, when extrapolating to mixtures of densely ionizing and sparsely ionizing radiation, ZRM systematically underestimates synergistic interactions (49–51) that enhance biological effectiveness. Specifically, if the densely ionizing component has small or zero *β*_*dense*_, which is common, then the quadratic cross-term 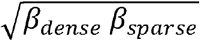 is small, or even vanishing, no matter how large the sparsely ionizing *β*_*sparse*_ term gets. This form of the cross-term therefore intrinsically limits the extent of ‘extra’ kill achievable by mixing, in contrast to empirical observations.

This systematic underestimation is reflective of a breakdown in the underlying assumptions built into ZRM and indicates the presence of additional mechanistic sources of synergy that are not captured. Specifically, ZRM assumes independent formation of sublethal lesions, random pairing, and identical interaction efficiencies across types (44,45,47). These conditions breakdown for combinations of highly disparate radiation qualities, as the densely ionizing radiation alters spatial colocalization and repair dynamics in ways that increase the efficiency of mixed quality interactions beyond a simple geometric mean.

Various approaches for going beyond the limitations of the ZRM have been proposed over the years (43,52). In general, however, these corrections still retain a feeling of being ad-hoc and arbitrary, with no clear or intuitive mechanistic basis that is universally accepted. This lack of a firm etiological foundation limits model interpretability, transparency, and trustworthiness, resulting in barriers to more widespread clinical adoption. A noteworthy recent example of such an approach is a proposed generalized BED formula based on a modified Zaider-Rossi model (mZRM (48)), where the combination of different quality radiations was modeled by taking the arithmetic mean of the quadratic coefficients. The authors reported improved fits to the existing data in the Particle Irradiation Data Ensemble (PIDE (53)). However, in the words of the authors themselves, *“The mZRM model is not based on a theoretical and classical biophysical mechanism of cell inactivation. This approach contrasts the direct-effect-mechanism bioeffect models such as the LEMs (local effect models), which relate DNA DSB distributions to local energy deposition*.*”*

### B. The MEDRAS Model

The Mechanistic DNA Repair and Survival (MEDRAS) model, developed by McMahon and colleagues, is a computational framework that simulates how cells repair radiation-induced DNA damage to predict biological outcomes like mutation and cell death. It takes a spatiotemporal distribution of DNA damage, particularly double strand breaks (DSBs), as input. This distribution may be the output of a detailed track structure simulation such as TOPAS-nBio. Alternatively, it can be generated using simpler analytic descriptions of track distribution, energy deposition, and resulting damage yield. This damage distribution is then fed into a model of the kinetics and fidelity of the major DSB repair pathways, including homologous recombination (HR), non-homologous end rejoining (NHEJ) and microhomology-mediated end rejoining (MMEJ). From this simulation, the model extracts various outcomes, including the overall final yield of chromosomal aberrations and the probability of cell death. Below, we describe each of these processes more explicitly.

The DSB distribution is generated starting from an ‘exposure’ object that characterizes the track structure of the radiation. The quality of a type of radiation is specified by the combination of its linear energy transfer *L* and atomic number Z, such that we may construct an arbitrary 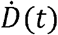 as a superposition 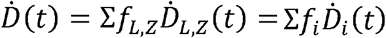. The quality-specific consequences of 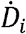 enter MEDRAS in one of two general ways: 1) modifying total and relative yields of simple vs. complex DSBs, which couple to downstream DNA damage response in distinct ways, and 2) modifying the statistics of DSB clustering and the relative magnitudes of inter-track vs. intra-track interactions.

DSBs in MEDRAS come in two groups. Simple DSBs, more common for sparsely ionizing radiation, have clean breaks with few modifications, are relatively easy to repair, and are less susceptible to early apoptosis. Complex DSBs, more commonly found in densely ionizing radiation, are accompanied by additional nearby damage that complicates repair. Simple and complex DSBs thus have different relative affinities for the various DSB repair pathways.

We can codify this information by assigning respective yield rates *Y*_*si*_, *Y*_*ci*_ for simple and complex DSBs for a given radiation quality *i*. Subsequently, we may assign simple and complex DSBs distinct probabilities of early cell death 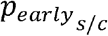 without attempted repair, and for those DSBs where repair is attempted, distinct probabilities that the attempt is made by the each of the three various pathways 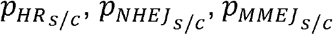, each with respective rates of repair *λ*_*HR*_, *λ*_*NHEJ*_, *λ*_*MMEJ*_.

In addition to overall yield and complexity distribution, radiation qualities also differ in how spatially clustered their DSBs are. Individual radiation tracks will traverse through a random cross section of the nucleus and generate DSBs at various points within that cross section. In the limit that the ionization density is extremely sparse, the probability that any given track generates more than one DSB is negligible. Thus, for many independent tracks, the resulting ‘inter-track’ distribution of DSBs is essentially spatially uncorrelated. However, for more densely ionizing radiation, if we randomly select a DSB, there is an increased chance that there will be additional nearby DSBs that originated from the same single-track cross section.

We can quantify this short-range clustering by borrowing from statistical mechanics (54) the concept of a pair correlation function *g*(*r*), which measures how the density of nearby DSBs varies as a function of distance from a reference DSB. In the absence of significant intra-track clustering we have *g*(*r*) = 1, in analogy to an ideal gas of point particles. Different radiation qualities, then, can be characterized by how much they deviate from 1 due to the details of their track structure 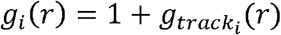, and the *g*(*r*) of a mixed beam can be constructed as the average of its constituents.

With this setup, we may proceed to calculate the overall probability of cell survival. MEDRAS incorporates three routes for cell death: early apoptotic cell death, mitotic catastrophe due to the fixation of unrepaired DSBs, and mitotic catastrophe due to lethal chromosomal aberrations arising from DSB misrepair. The first two contributions are relatively straightforward to model. We set *t* = 0 as the start of the exposure, and suppose that the cell is in a ‘pre-checkpoint’ state for a time *t*_*r*_. Then, we can integrate the effective *p*_*early*_ of all unrepaired DSBs from *t* = 0 to *t*_*r*_ to get the cumulative rate of early cell death. Likewise, if we integrate the system of differential equations for the three repair pathways, we can solve for the time-dependent number of unrepaired DSBs, which we then evaluate at t_r_ to get the first mitotic catastrophe term.

The third contribution, mitotic catastrophe arising from lethal chromosomal aberrations, requires additional analysis and information. We again integrate the system of differential equations for three repair pathways, but this time we need the rate of binary misrepair for spatially separated DSBs relative to the rate of correct local repair. We also need to calculate the likelihood that the resulting chromosomal aberration is lethal. These quantities are controlled by a combination of cell-specific parameters, including (25) the spatial range of misrepair *σ*, the volume of the nucleus *V*_*nuc*_, the volume of a chromosomal domain *V*_*dom*_ and the lethal threshold for gene deletion *L*_*D*_.

## III. Methods: MBAM Algorithm and Application to MEDRAS

In this section, we introduce the technical details and building blocks of MBAM. To keep the discussion concrete, we consider the specific radiobiological problem of predicting the cell survival curve *S*(*D*) with MEDRAS. For this paper, we restrict ourselves to the case of an *acute* exposure, with some total dose *D* delivered instantaneously at *t* = 0, distributed over *N* radiation types

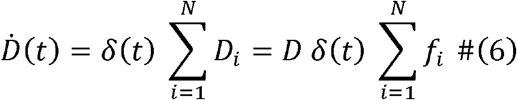

Formally, the MBAM algorithm works (37,40) by approximating high-dimensional models with simplified representations on the “boundaries” of a model’s parameter space, or model manifold. The manifold represents all possible outputs the model can produce given different parameter values. To reduce this model, one must calculate eigenvalues and eigenvectors of the Fisher Information Matrix (FIM) to find the least relevant parameter combinations. This guides the process to a “boundary” on the parameter space manifold, where a simplified model is constructed by removing the identified parameter combination. This new, simplified model can be recalibrated to match the behavior of the original model. The process can be repeated to further to yield a hierarchy of reduced models that maintain predictive fidelity, while systematically removing degrees of freedom until the desired level of reduction is achieved.

Explicitly, 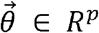 is the full parameter vector for the complex model, and 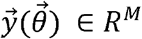 is some vector of model predictions for quantities of interest. In the context of our radiobiological application, 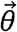 is the set of exposure-specific and cell-specific MEDRAS parameters while 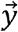 is the measured cell survival *S* for *M* non-zero absorbed doses. The parameter vector 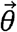, for an N-component radiation mixture, includes:

- *4N* radiation-specific exposure parameters (*f,Y*_*s*_, *Y*_*c*_,*g*_*track*_)_*i*_ for i = 1, …, N
- 8 complexity-dependent probabilities (*p*_*early*_, *p*_*HR*_, *p*_*NHEJ*_, *p*_*MMEJ*_)_*s/c*_
- 3 repair rates (*λ*_*HR*,_*λ*_*NHEJ*,_ *λ*_*MMEJ*_)
- 5 remaining cell-specific parameters (*t*_*r*_, *σ, V*_*nuc*_,*V*_*dom*_,*L*_*D*_)

We would like to map an arbitrary initial parameter vector 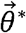 in the high-dimensional p = 4N + 16 parameter space of MEDRAS to a smaller set of effective parameters that are more easy to associate with the phenomenological LQ paradigm. To do this, start by computing the Jacobian matrix 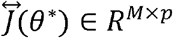

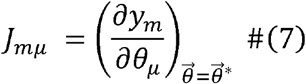

From this, the FIM *g* (*θ*^*\**^) ∈ *R*^*p*×*p*^ is obtained as

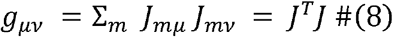

We now solve the eigenproblem 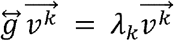, and order the eigenvalues *λ*_1_ ≤ *λ*_2_ ≤ … ≤ *λ*_p_ The eigenvector 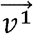 corresponding to the smallest eigenvalue defines the softest (sloppiest) local direction in parameter space, which can be interpreted as the collective variation of MEDRAS parameters that least changes predictions for the cell survival curve *S*(*D*).

MBAM moves along a geodesic on the model manifold in this sloppiest direction, and follows it to a boundary in parameter space. If we define a geodesic coordinate *τ*, we may say our parameter vector has initial position 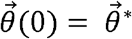 and initial velocity 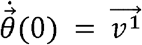. We thus integrate to an updated point in parameter space 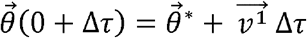, and from this new point we iterate and repeat the process to trace out the full geodesic curve 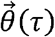 starting from our initial 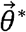.

As *τ* → ∞, the sloppiest eigenvalue *λ*_1_ → 0. This can be interpreted as the geodesic having travelled to a limiting region in parameter space where the sloppiest direction in parameter space is *practically unidentifiable* with respect to predictions for the quantities of interest. What this means is that we may reduce the number of independent fit parameters in our model by making some kind of *interpretable approximation*. For example, the MMEJ pathway is known to be much slower than either the HR or NHEJ pathways, so a realistic initial point in MEDRAS parameter space has a good chance of moving along a geodesic to a region of parameter space where *λ*_*HR*_, *λ*_*NHEJ*_ ≫ *λ*_*MMEJ*_, such that, effectively, we can take *λ*_*MMEJ*_ ≈0 and remove it from the model.

The reduced set of parameters 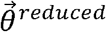 from this round of MBAM can then be fit to predictions for the quantities of interest based on the original point in parameter space 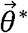, to get an effective ‘mapping’ 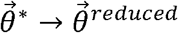. The next round of MBAM would then remove the next-sloppiest parameters in the model, for example, by setting various probabilities to 0 or 1 or having 0 ≪ *t*_*r*_ → ∞. In this way, the dimensionality of the parameter vector successively shrinks as the least important combinations of parameters are ‘interpretably approximated’ out.

It is worth pointing out two situations that sometimes arise. Firstly, if the sloppiest eigenvalue *λ*_*1*_ = 0, then the corresponding sloppy eigenvector becomes *not just practically*, but *structurally, unidentifiable*. It becomes impossible, even in principle, to uniquely determine its value using the quantities of interest. In these cases, the effective parameters of the reduced model 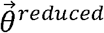 have an exact closed-form expression in terms of the original parameters of the full model 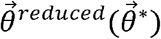 Secondly, the MBAM reduction can sometimes give a ‘multiple-for-one’ deal, where approximating out one parameter automatically removes one or more additional parameters. For example, if the yield of complex damage is negligible for all the component exposures *Y*_*ci*_ ≈ 0, then it is easy to see that any downstream model predictions in the reduced model will also be independent of the four ‘complex-damage-specific’ probabilities.

## IV. Results: Bridging MEDRAS with LQ Phenomenology

In this section, we present the results of our application of MBAM to MEDRAS. In subsection IV.A. we present the results of early stage of model reduction, where the full MEDRAS model for a general mixed-quality acute exposure is found to be *structurally indistinguishable* from a reduced three-parameter model for an effective uniform quality exposure, which we call MEDRAS-LPL. In subsection IV.B., we further analyze MEDRAS-LPL, identifying two distinct manifold boundaries in MEDRAS-LPL parameter space, corresponding to two different limiting approximations that enforce ‘boundary conditions’ on the corresponding effective LQ parameters. Finally, in section IV.C., we discuss the implications of these results for model-based inference, prediction, and optimization of mixed-quality radiation response.

### A. MEDRAS is Structurally Indistinguishable from MEDRAS-LPL

Initial iterations of MBAM indicate that, when predicting *S*(*D*) for acute exposures using MEDRAS, all but three of the FIM eigenvalues are *exactly zero!* As described in the previous section, the presence of eigenvalues equal to zero indicates structural unidentifiability and allows for an explicit mapping of the original parameters of the complex model to the effective parameters of the reduced model. The mathematical form of this reduced model for MEDRAS, as seen in **Figure 2**, bears a striking resemblance to the Lethal-Potentially Lethal (LPL) model of Curtis (55).

**Figure 2:**
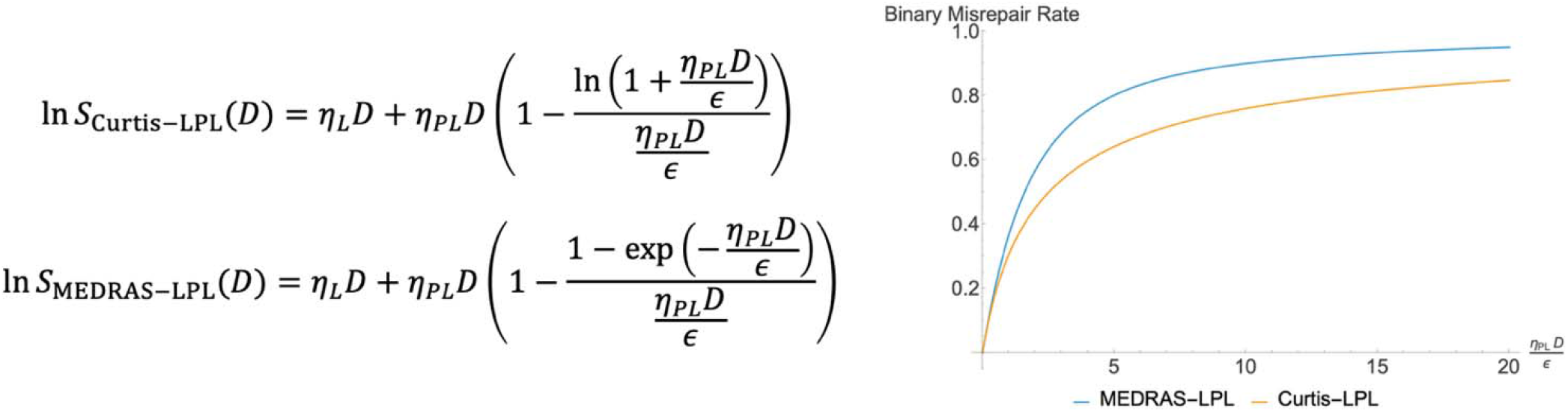
The first level of MBAM reduction on MEDRAS yields a reduced MEDRAS-LPL model, similar to the original LPL model of Curtis but with a different nonlinearity.

The LPL model describes how cell death occurs after radiation exposure by differentiating between two types of lesions: lethal lesions, produced at a rate per unit dose *η*_*L*_, and potentially lethal lesions, produced at a rate per unit dose *η*_*PL*_ which can undergo either correct repair or binary misrepair. The relative rate of misrepair *∈* following an acute single fraction exposure scales nonlinearly. At sufficiently low absorbed doses, *∈* is linearly proportional to the total amount of potentially lethal lesions. However, as the dose increases, the buildup of damage saturates the cellular repair machinery, with *∈* →1 in the limit that *D* → ∞ and repair becomes completely inefficient.

As shown in **Figure 3**, the result of reducing MEDRAS is an effective three-parameter model that is almost identical to the LPL. Accordingly, we may refer to this reduced model as MEDRAS-LPL. The only minor difference between mEdRAS-LPL and the original LPL is in the nonlinear scaling of repair saturation, which is faster for MEDRAS-LPL. However, in contrast to the generic LPL model, the MEDRAS-LPL effective parameters can be decomposed into mechanistically interpretable contributions, which becomes useful when seeking to understand how various biological and physical ‘control knob’ parameters may modulate biological response.

**Figure 3:**
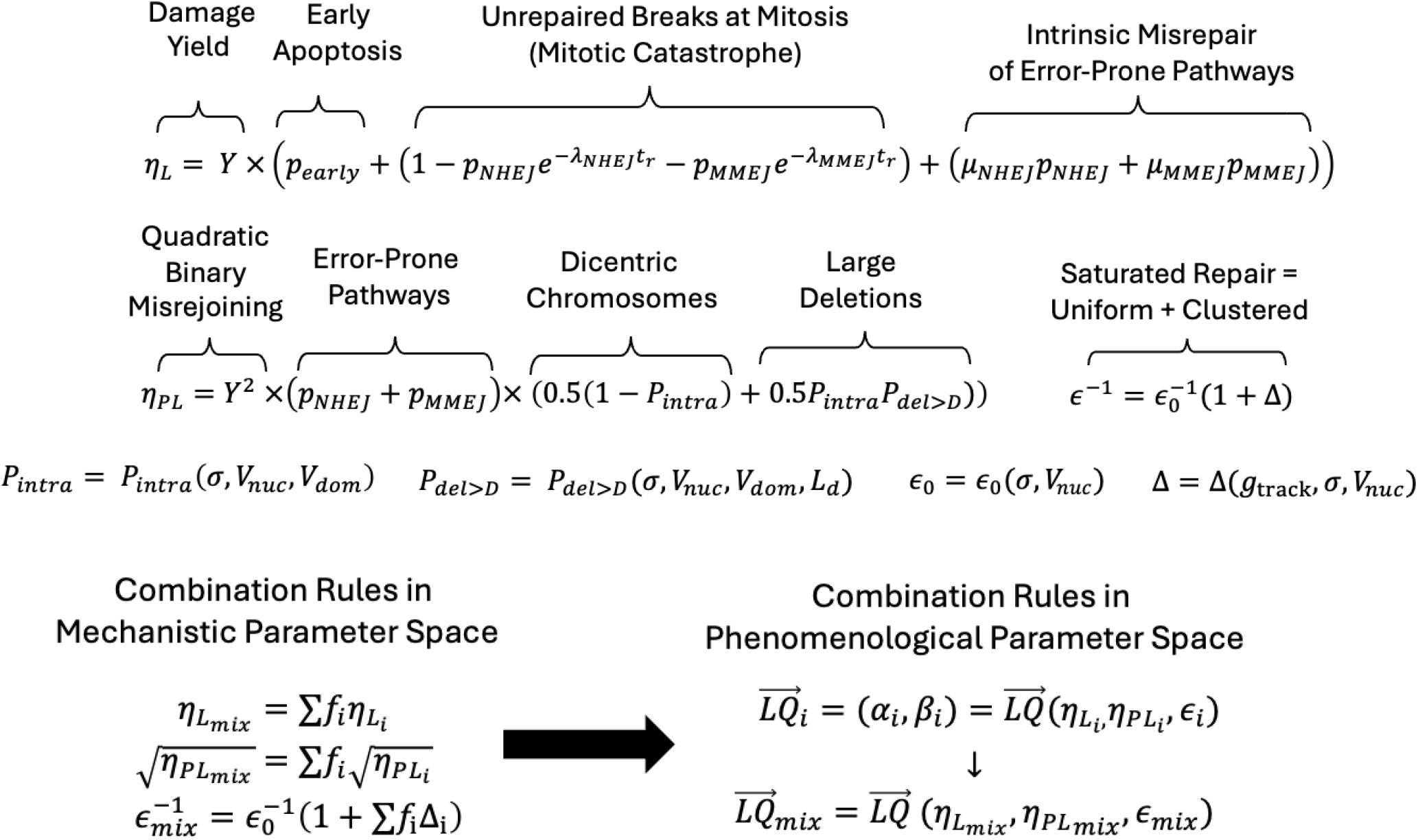
The derivation of MEDRAS-LPL from MEDRAS allows for an interpretable and systematic breakdown of how the various mechanistic processes enter into the effective parameters (top), leading to elegant and interpretable combination rules in MEDRAS-LPL parameter space that provide a natural strategy for identify the corresponding combination rules in LQ space (bottom).

In the context of mixed-quality response, the control knobs of interest are the relative fractions *f*_*i*_ of the different component qualities in the mixture. The explicit decomposition of the effective parameters in terms of the original parameters makes it straightforward to see that the effective *η*_*L*_ and *η*_*PL*_ of the mixed exposure are the arithmetic and geometric means of the components, respectively, in line with our intuitive expectations based on the theory of dual radiation action and the ZRM paradigm.

However, we additionally see now that the effective relative misrepair rate *e also* combines in an intuitive and interpretable manner based on the underlying MEDRAS mechanisms. Specifically, we exploit the fact that *g*(*r*) = 1 + *g*_*track*_ (*r*) to write *∈* = *∈*_0_(l + Δ). Here, *∈*_0_ corresponds to *g*(*r*) = 1, and is the minimum possible value for the case of negligible intra-track effects, with misrepair events being dominated by uniform, and randomly distributed inter-track interactions. Meanwhile, Δ is the excess misrepair arising from intra-track effects due to non-random clustering that increases the interaction cross section for binary misrepair. Written this way, it is straightforward to see that *∈*_0_ is a cell-specific quantity that is independent of radiation quality, such that any quality-dependence comes entirely through Δ, which for a mixed beam is just the usual arithmetic mean of its components.

### B. Mapping MEDRAS-LPL to LQ Phenomenology

The previous results demonstrate that we have reduced the problem of predicting LQ radiosensitivity for an arbitrary mixed exposure to the problem of mapping some general (*η*_*L*_*η*_;*PL*_*∈*) to a corresponding set of (*α,β*) parameters. To this end, in this subsection we present results for the final iteration of MBAM applied to MEDRAS-LPL. Here, unlike for the generic MEDRAS model that we started with, the sloppiest eigenvalue is small, but not exactly equal to zero, which prevents us from writing explicit analytic closed-form expression for additional reductions.

However, the MBAM calculations here still lead to a surprising ‘two-for-one’ result where the boundaries of the model manifold reduce the three-parameter model to an effective one-parameter description. A vector field plot of the sloppy eigenvector, illustrated in **Figure 4**, indicates the presence of two different manifold boundaries, corresponding to two qualitatively different interpretable limiting approximations in parameter space.

**Figure 4:**
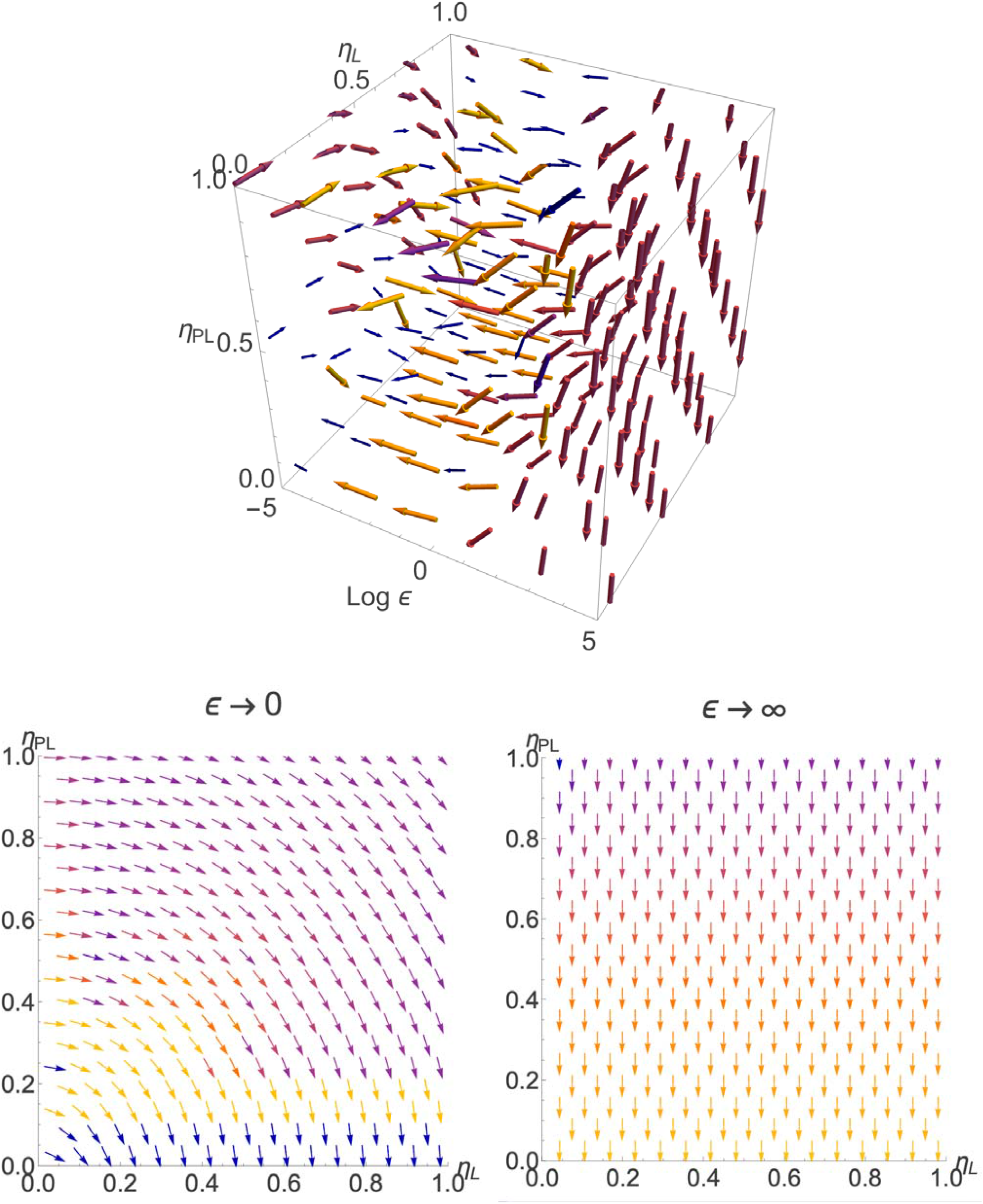
The final round of MBAM on the MEDRAS-LPL model (top) identifies a vector field of sloppy geodesics that flow to one of two manifold boundaries, corresponding to one of two interpretable approximations (bottom), with *∈* → 0 corresponding to densely ionizing radiation and *∈* → ∞ to sparsely ionizing radiation.

As *∈* → ∞, the curves of the geodesic all tend towards *η*_*PL*_ →0, indicating the limit of negligible binary misrepair, such that only directly lethal damage is significant. In other words, for a MEDRAS-LPL model with very large *∈*, the parameter *η*_*PL*_ *automatically* becomes structurally unidentifiable, since if there is no possibility of misrepair, any potentially lethal damage automatically becomes nonlethal, and the MEDRAS-LPL model reduces exactly to a linear model with *α* = *η*_*L*_,

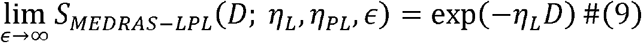

On the other hand, as *∈* →0, the curves of the geodesic all tend towards a constant value of *η*_*L*_ + *η*_*PL*_, indicating the opposite limit of negligible correct repair, such that all potentially lethal damage becomes lethal, in effect getting absorbed into the directly lethal damage term. This limit, too, is a linear model, but with *a –η*_*L*_ + *η* _*PL*_

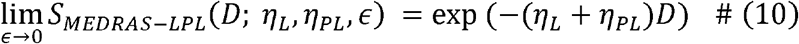

As another way of interpreting these boundaries, we note that the limiting forms of *a* for *∈*→ ∞ and *∈*→0 line up with previously derived results from the Curtis LPL model for sparsely and densely ionizing radiation, respectively (55). For the *∈*→ ∞, the linear coefficient may be thought of as the first-order term in a Taylor series expansion of a ZRM dual radiation action model. Meanwhile, for the *∈*→ 0 side, it is more appropriate to think of the system as approaching a target theoretic ‘single-target-single-hit’ description, with the linear coefficient akin to the average number of hits per unit dose.

Note that there is no explicit quadratic term *β* at either boundary. The implication of this result is that, starting from MEDRAS-LPL, the most universal interpretation of the LQ model is in terms of *phenomenological* curve fitting: a given (*η*_*L*_,*η*_*PL*_, *∈*) generates a simulated survival curve *S*_*MEDRAS*_*−*_*LPL*_(*D; η*_*L*_,*η*_*PL*_, *∈*). If we fit this simulated data to an LQ model with fitting parameters 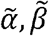 the ‘true’ LQ coefficients are just those that minimize the sum of squares loss *L* over all simulated doses *D*_*i*_,.

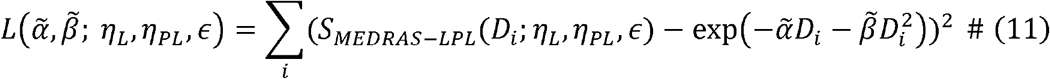

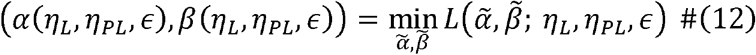

As mentioned previously, the explicit form of the *α* and *β* mapping as a function of (*η*_*L*_, *η*_*PL*_, *∈*) is not expressible in closed form for general MEDRAS-LPL parameters, since the sloppy eigenvalue is not exactly equal to zero. However, the MBAM limits do place formally exact *boundary conditions* on the form of the mapping

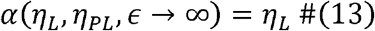

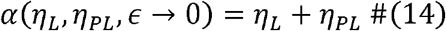

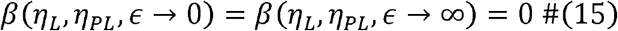

Away from the manifold boundaries, the effective *α, β* mapping must be determined by direct nonlinear least-squares fitting. **Figure 5** displays the results of this mapping for a series of intermediate values of *∈*. The best-fit linear parameter *a* is universally observed to be a linear function *η*_*L*_ and *η*_*PL*_, with the relative slope of this line varying with *∈*. Meanwhile, the best-fit quadratic parameter *β* is seen to be independent of *η*_*L*_ regardless of *∈*. The extreme ends of *∈* further enforce ‘hump-like’ behavior, such that the *β*(*η*_*PL*_, *∈*) curve initially rises, then beyond some crossover *∈*, must drop again to satisfy the requirement that *β* ⟶ 0 at the manifold boundaries. Framed another way, the MBAM limits imply that there must be an ‘inflection surface’ of local maxima in *β*(*η*_*PL*_, *∈*).

**Figure 5:**
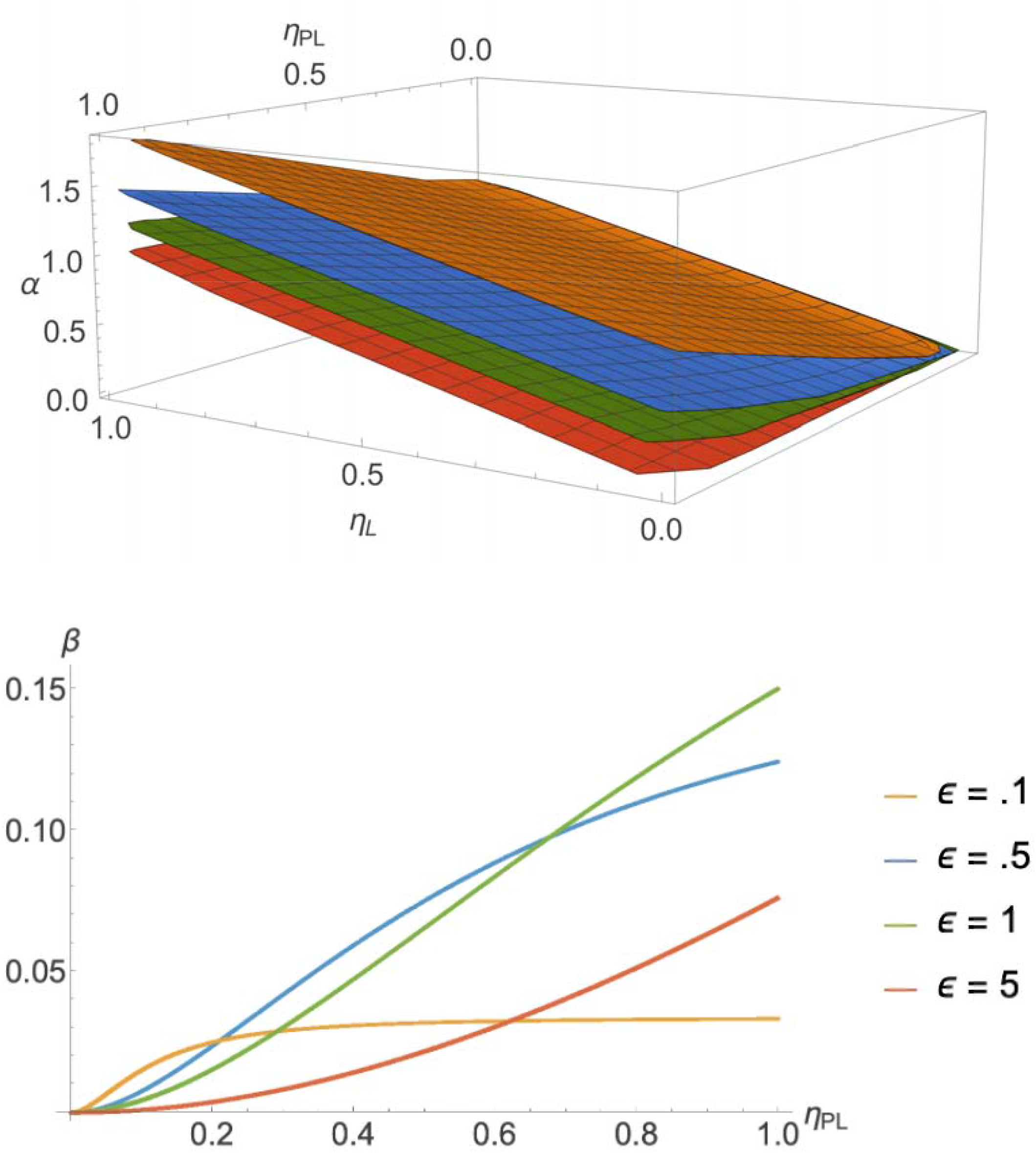
Mapping MEDRAS-LPL parameter space to LQ parameters shows that the effective linear parameter α is a linear combination of η_L_ and η_PL_ with e-dependent slope (top). Meanwhile, the quadratic parameter β rises and falls with ∈, in line with the zero boundary conditions resulting from the MBAM limits.

We do note, however, that for MEDRAS-LPL parameters that are close to the *∈* ⟶ ∞ boundary, the form of *β*(*η*_*PL*_, *∈*) becomes progressively closer to a directly proportional linear dependence, as is expected from the theory of dual radiation action. Thus we see that, while our operational definition of *a* and *β* in terms of goodness-of-fit is more universal, it passes the basic ‘sanity test’ of giving results equivalent to the ZRM paradigm for sparsely ionizing radiation.

### C. Implications for Mixed Quality Response: Optimizing Synergistic Interactions

Putting the results of the previous two subsections together, we arrive at an elegant and natural formalism to mixed-quality radiation response prediction. Significantly, this approach goes beyond being simply empirical interpolation of measured data, but in fact extrapolates to generate truly novel, experimentally testable hypotheses.

To demonstrate this capability, we may consider a hypothetical mixture of two radiation qualities. One type of radiation has MEDRAS-LPL parameters close to the sparsely ionizing *∈* → ∞ boundary, while the other has parameters close to the densely ionizing *∈* → 0 boundary. We may map an arbitrary fraction composition of these two qualities onto those of an ‘equivalent uniform quality’ 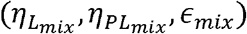 using the results of section IV.A, particularly, the combination rule reported in **Figure 3**.

From the results of section IV.B., we see that it is straightforward to map this effective 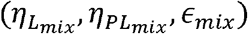 onto corresponding effective 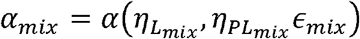 and 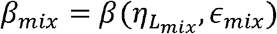 LQ coefficients. From these, we can generate the corresponding effective LQ survival curve for the mixture. The results of this procedure are illustrated in **Figure 6** for different compositions of sparse and dense radiation. The calculations of the model clearly point to the enhancement of cell kill beyond the predictions of the ZRM paradigm, in line with experimental and clinical observation.

**Figure 6:**
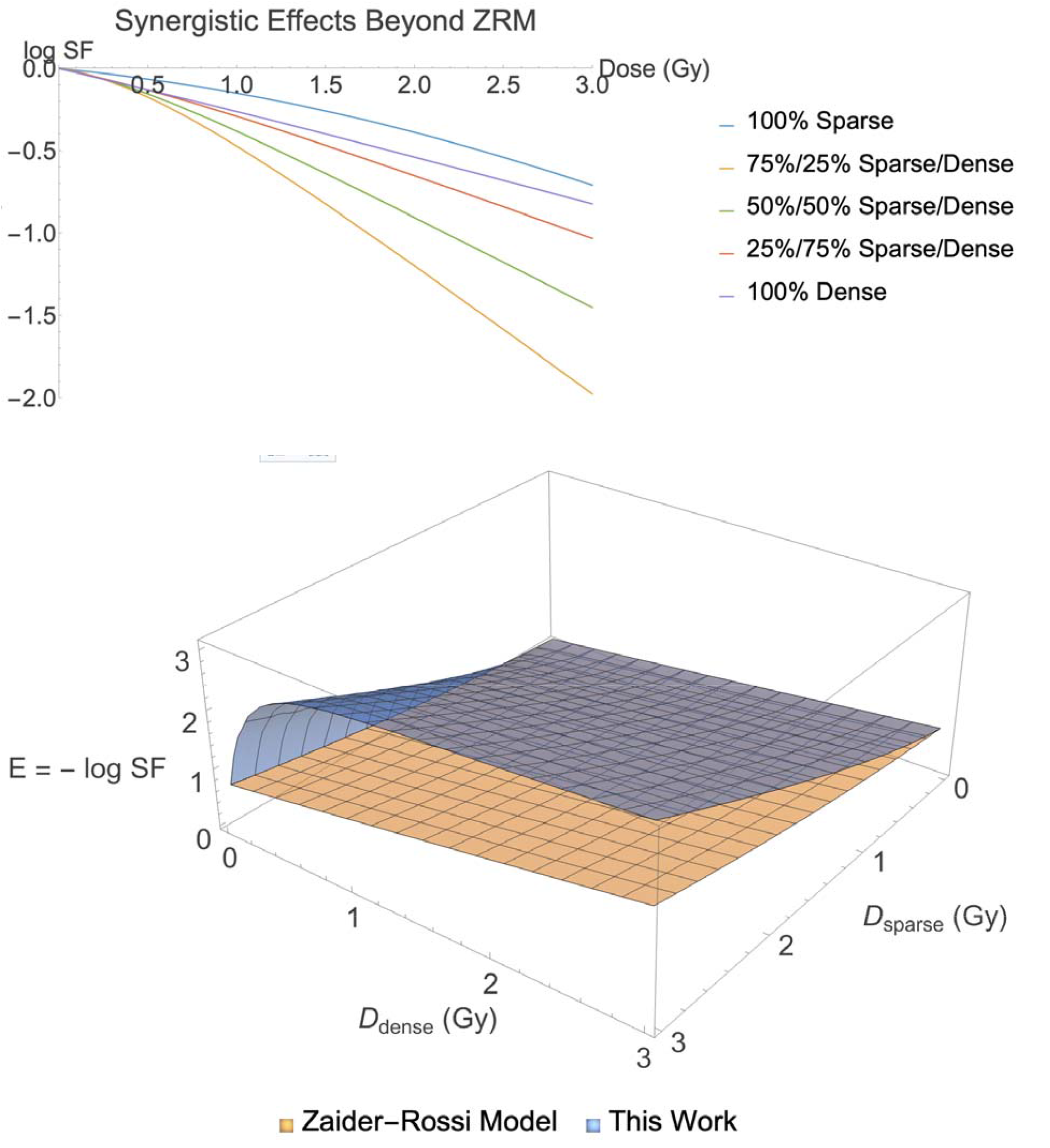
The results of our model predict synergistic enhancement of S(D) for dense and sparse radiation beyond the calculations ZRM model, with the ‘peak’ cytotoxicity occurring at some intermediate composition that can be systematically estimated and zoned in on using the model-informed approach developed in this study.

Furthermore, our model also predicts as a corollary an ‘optimal’ composition of sparse and dense radiation that maximizes cytotoxicity. This result has a natural radiobiological interpretation based on the mechanistic underpinnings we have derived it from. In the example shown, sparsely ionizing radiation creates more damage per unit dose, but the fraction of damage that is directly lethal is low, as most of it is unclustered and thus has a smaller low cross section for binary misrepair, leading to relatively high repair efficiency. On the other hand, the example densely ionizing radiation creates less total damage per unit dose, but what damage it does create is more clustered, and as a result, more rapidly saturates the repair processes. When combining the two types of radiation, this creates a tradeoff between total damage and overall repair efficiency. At some sweet-spot composition, this tradeoff is optimized, leading to maximal cell kill.

## V. Summary and Future Work

In summary, we have demonstrated that systematic model reduction serves as a general strategy for deriving parsimonious, mechanism-based reduced-order descriptors of biological effectiveness from detailed mechanistic radiobiological knowledge encoded in complex multi-parameter models. Notably, our approach goes beyond empirical interpolation of ‘black-box’ measurements of LQ parameters under different conditions, to extrapolative generation of new hypotheses. Such hypotheses can be used to guide the design of new experiments, to enable truly predictive, model-informed optimization in radiobiology.

The results presented here open many avenues for future exploration. Most immediately, the prediction of an optimal intermediate mixture of sparse and dense ionizing radiation that maximizes cytotoxicity, along with our model-based strategy for systematically identifying this optimal mixture, can be tested experimentally. Beyond this, however, there remains much room for further work integrating ever more detailed, state-of-the-art mechanistic Monte Carlo radiobiological simulations, such as Geant4-DNA, with the MBAM formalism. To this end, an important methodological extension to prioritize will be the extension of the FIM calculation beyond deterministic ODE models to more general stochastic Monte Carlo calculations. In addition, it will be worthwhile to investigate how to integrate this information geometric analysis with surrogate neural network models (56) to accommodate situations where we may have only partial knowledge of the underlying mechanisms and parametric processes.

## REFERENCES

1. McMahon SJ. The linear quadratic model: usage, interpretation and challenges. Phys Med Biol. 2018 Dec 19;64(1):01TR01.

2. C. Joiner M, Van Der Koge J.LA. Basic Clinical Radiobiology [Internet]. 6th ed. Boca Raton: CRC Press; 2024 [cited 2025 Jun 11]. Available from: https://www.taylorfrancis.com/books/9781003278337

3. Hall EJ. Radiobiology for the radiologist. 1988 [cited 2021 Oct 1]; Available from: http://inis.iaea.org/Search/search.aspx?orig_q=RN:21040148

4. McMahon SJ, McNamara AL, Schuemann J, Paganetti H, Prise KM. A general mechanistic model enables predictions of the biological effectiveness of different qualities of radiation. Sci Rep. 2017 Sep 7;7(1):10790.

5. Fowler JF. The linear-quadratic formula and progress in fractionated radiotherapy. Br J Radiol. 1989 Aug 1;62(740):679–94.

6. Fowler JF. 21 years of Biologically Effective Dose. Br J Radiol. 2010 Jul;83(991):554–68.

7. Jeggo P, Lavin MF. Cellular radiosensitivity: How much better do we understand it? Int J Radiat Biol. 2009 Dec;85(12):1061–81.

8. Fertil B, Malaise EP. Inherent cellular radiosensitivity as a basic concept for human tumor radiotherapy. Int J Radiat Oncol. 1981 May;7(5):621–9.

9. Gray LH, Conger AD, Ebert M, Hornsey S, Scott OCA. The Concentration of Oxygen Dissolved in Tissues at the Time of Irradiation as a Factor in Radiotherapy. Br J Radiol. 1953 Dec;26(312):638–48.

10. Tannock IF. Oxygen diffusion and the distribution of cellular radiosensitivity in tumours. Br J Radiol. 1972 Jul;45(535):515–24.

11. Kirkpatrick JP, Meyer JJ, Marks LB. The Linear-Quadratic Model Is Inappropriate to Model High Dose per Fraction Effects in Radiosurgery. Semin Radiat Oncol. 2008 Oct;18(4):240–3.

12. Guerrero M, Li XA. Extending the linear-quadratic model for large fraction doses pertinent to stereotactic radiotherapy. Phys Med Biol. 2004 Oct 21;49(20):4825–35.

13. Lea DE, Catcheside DG. The mechanism of the induction by radiation of chromosome aberrations inTradescantia. J Genet. 1942 Dec;44(2-3):216–45.

14. Franken NAP, Oei AL, Kok HP, Rodermond HM, Sminia P, Crezee J, et al. Cell survival and radiosensitisation: Modulation of the linear and quadratic parameters of the LQ model. Int J Oncol. 2013 May;42(5):1501–15.

15. Van Leeuwen CM, Oei AL, Crezee J, Bel A, Franken NAP, Stalpers DA, et al. The alfa and beta of tumours: a review of parameters of the linear-quadratic model, derived from clinical radiotherapy studies. Radiat Oncol. 2018 Dec;13(1):96.

16. Hall EJ. Radiation Dose-Rate: A Factor of Importance in Radiobiology and Radiotherapy. Br J Radiol. 1972 Feb;45(530):81–97.

17. Gordon Steel G, Down JD, Peacock JH, Stephens TC. Dose-rate effects and the repair of radiation damage. Radiother Oncol. 1986 Jan;5(4):321–31.

18. Azevedo TA, Abrantes AM, Carvalho J. Radiobiological Modeling with Monte Carlo Tools - Simulating Cellular Responses to Ionizing Radiation. Technol Cancer Res Treat. 2025 Jun;24:15330338251350909.

19. McMahon SJ, Prise KM. Mechanistic Modelling of Radiation Responses. Cancers. 2019 Feb 10;11(2):205.

20. Gardner LL, Thompson SJ, O’Connor JD, McMahon SJ. Modelling radiobiology. Phys Med Biol. 2024 Sep 21;69(18):18TR01.

21. Chatzipapas KP, Papadimitroulas P, Emfietzoglou D, Kalospyros SA, Hada M, Georgakilas AG, et al. Ionizing Radiation and Complex DNA Damage: Quantifying the Radiobiological Damage Using Monte Carlo Simulations. Cancers. 2020 Mar 26;12(4):799.

22. Incerti S, Baldacchino G, Bernal M, Capra R, Champion C, Francis Z, et al. THE GEANT4-DNA PROJECT. Int J Model Simul Sci Comput. 2010 Jun;01(02):157–78.

23. Schuemann J, McNamara AL, Ramos-Méndez J, Perl J, Held KD, Paganetti H, et al. TOPAS-nBio: An Extension to the TOPAS Simulation Toolkit for Cellular and Sub-cellular Radiobiology. Radiat Res. 2018 Jan 4;191(2):125.

24. McMahon SJ, Schuemann J, Paganetti H, Prise KM. Mechanistic Modelling of DNA Repair and Cellular Survival Following Radiation-Induced DNA Damage. Sci Rep. 2016 Sep 14;6(1):33290.

25. McMahon SJ, Prise KM. A Mechanistic DNA Repair and Survival Model (Medras): Applications to Intrinsic Radiosensitivity, Relative Biological Effectiveness and Dose-Rate. Front Oncol. 2021 Jun 29;11:689112.

26. Lim A, Andriotty M, Yusufaly T, Agasthya G, Lee B, Wang C. A fast Monte Carlo cell-by-cell simulation for radiobiological effects in targeted radionuclide therapy using pre-calculated single-particle track standard DNA damage data. Front Nucl Med. 2023 Dec 6;3:1284558.

27. Ramos-Méndez J, Perl J, Schuemann J, McNamara A, Paganetti H, Faddegon B. Monte Carlo simulation of chemistry following radiolysis with TOPAS-nBio. Phys Med Biol. 2018 May 17;63(10):105014.

28. Ramos-Méndez J, Domínguez-Kondo N, Schuemann J, McNamara A, Moreno-Barbosa E, Faddegon B. LET-Dependent Intertrack Yields in Proton Irradiation at Ultra-High Dose Rates Relevant for FLASH Therapy. Radiat Res [Internet]. 2020 Aug 28 [cited 2026 Jan 20];194(4). Available from: https://bioone.org/journals/radiation-research/volume-194/issue-4/RADE-20-00084.1/LET-Dependent-lntertrack-Yields-in-Proton-lrradiation-at-Ultra-High/10.1667/RADE-20-00084.1.full

29. Zhu H, McNamara AL, McMahon SJ, Ramos-Mendez J, Henthorn NT, Faddegon B, et al. Cellular Response to Proton Irradiation: A Simulation Study with TOPAS-nBio. Radiat Res. 2020 May 13;194(1):9.

30. Zhu H, McNamara AL, Ramos-Mendez J, McMahon SJ, Henthorn NT, Faddegon B, et al. A parameter sensitivity study for simulating DNA damage after proton irradiation using TOPAS-nBio. Phys Med Biol. 2020 Apr 21;65(8):085015.

31. Lai Y, Tsai M, Tian Z, Qin N, Yan C, Hung S, et al. A new open-source GPU-based microscopic Monte Carlo simulation tool for the calculations of DNA damages caused by ionizing radiation — Part II: sensitivity and uncertainty analysis. Med Phys. 2020 Apr;47(4):1971–82.

32. Thompson SJ, Prise KM, McMahon SJ. Monte Carlo damage models of different complexity levels predict similar trends in radiation induced DNA damage. Phys Med Biol. 2024 Nov 7;69(21):215035.

33. Snowden TJ, Van Der Graaf PH, Tindall MJ. Methods of Model Reduction for Large-Scale Biological Systems: A Survey of Current Methods and Trends. Bull Math Biol. 2017 Jul;79(7):1449–86.

34. Evangelou N, Wichrowski NJ, Kevrekidis GA, Dietrich F, Kooshkbaghi M, McFann S, et al. On the parameter combinations that matter and on those that do not: data-driven studies of parameter (non)identifiability. Nelson KE, editor. PNAS Nexus. 2022 Sep 1;1(4):pgac154.

35. Sinitsyn NA, Hengartner N, Nemenman I. Adiabatic coarse-graining and simulations of stochastic biochemical networks. Proc Natl Acad Sci. 2009 Jun 30;106(26):10546–51.

36. Radulescu O, Gorban AN, Zinovyev A, Noel V. Reduction of dynamical biochemical reactions networks in computational biology. Front Genet [Internet]. 2012 [cited 2026 Jan 20];3. Available from: http://journal.frontiersin.org/article/10.3389/fgene.2012.00131/abstract

37. Transtrum MK, Qiu P. Model Reduction by Manifold Boundaries. Phys Rev Lett. 2014 Aug 29;113(9):098701.

38. Gutenkunst RN, Waterfall JJ, Casey FP, Brown KS, Myers CR, Sethna JP. Universally Sloppy Parameter Sensitivities in Systems Biology Models. Arkin AP, editor. PLOS Comput Biol. 2007 Oct 5;3(10):e189.

39. Transtrum MK, Machta BB, Brown KS, Daniels BC, Myers CR, Sethna JP. Perspective: Sloppiness and emergent theories in physics, biology, and beyond. J Chem Phys. 2015 Jul 7;143(1):010901.

40. Quinn KN, Abbott MC, Transtrum MK, Machta BB, Sethna JP. Information geometry for multiparameter models: New perspectives on the origin of simplicity. 2021 [cited 2022 Mar 11]; Available from: https://arxiv.org/abs/2111.07176

41. White A, Tolman M, Thames HD, Withers HR, Mason KA, Transtrum MK. The Limitations of Model-Based Experimental Design and Parameter Estimation in Sloppy Systems. Csikász-Nagy A, editor. PLOS Comput Biol. 2016 Dec 6;12(12):e1005227.

42. DeTal N, Anderson CNK, Transtrum MK. A CRISP approach to QSP: XAI enabling fit-for-purpose models [Internet]. arXiv; 2025 [cited 2026 Jan 19]. Available from: https://arxiv.org/abs/2505.02750

43. Parousis-Paraskevas O, Gkikoudi A, Al-Qaaod A, Vasilopoulos SN, Manda G, Beinke C, et al. Combined Radiations: Biological Effects of Mixed Exposures Across the Radiation Spectrum. Biomolecules. 2025 Sep 5;15(9):1282.

44. Zaider M, Rossi HH. The synergistic effects of different radiations. Radiat Res. 1980 Sep;83(3):732–9.

45. Rossi HH, Zaider M. Microdosimetry and Its Applications [Internet]. Berlin, Heidelberg: Springer Berlin Heidelberg; 1996 [cited 2026 Jan 19]. Available from: http://link.springer.com/10.1007/978-3-642-85184-l

46. Kellerer AM, Rossi HH. A Generalized Formulation of Dual Radiation Action. 1978 Sep 1 [cited 2026 Jan 19]; Available from: https://radiation-research.kglmeridian.eom/view/journals/rare/75/3/article-p471.xml

47. Kellerer AM, Rossi HD. THEORY OF DUAL RADIATION ACTION. Curr Top Radiat Res Quart 8 No 2 85-1580ct 1972 [Internet]. 1971 Dec 31 [cited 2026 Jan 19]; Available from: https://www.osti.gov/biblio/4611340

48. Katugampola S, Hobbs RF, Howell RW. Generalized methods for predicting biological response to mixed radiation types and calculating equieffective doses (EQDX). Med Phys. 2024;51(1):637–49.

49. Ham DW, Song B, Gao J, Yu J, Sachs RK. Synergy Theory in Radiobiology. Radiat Res. 2017 Dec 29;189(3):225.

50. Suzuki S. The ‘Synergistic’ Action of Mixed Irradiation with High-LET and Low-LET Radiation. Radiat Res. 1994 May;138(2):297.

51. Lam GKY. The Survival Response of a Biological System to Mixed Radiations. Radiat Res. 1987 May;110(2):232.

52. Taleei R, Rahmanian S, Nikjoo H. Modelling Cellular Response to Ionizing Radiation: Mechanistic, Semi-Mechanistic, and Phenomenological Approaches - a Historical Perspective. Radiat Res [Internet]. 2024 Jun 25 [cited 2026 Jan 20];202(2). Available from: https://bioone.org/journals/radiation-research/volume-202/issue-2/RADE-24-00019.l/Modelling-Cellular-Response-to-lonizing-Radiation--Mechanistic-Semi-Mechanistic/10.1667/RADE-24-00019.1.full

53. Friedrich T, Pfuhl T, Scholz M. Update of the particle irradiation data ensemble (PIDE) for cell survival. J Radiat Res (Tokyo). 2021 Jul 10;62(4):645–55.

54. Chandler D. Introduction to Modern Statistical Mechanics. Oxford, New York: Oxford University Press; 1987. 288 p.

55. Curtis SB. Lethal and potentially lethal lesions induced by radiation--a unified repair model. Radiat Res. 1986 May;106(2):252–70.

56. Gherman IM, Abdallah ZS, Pang W, Gorochowski TE, Grierson CS, Marucci L. Bridging the gap between mechanistic biological models and machine learning surrogates. Schulz MH, editor. PLOS Comput Biol. 2023 Apr 20;19(4):el010988.

